# The Crystal Structure of the Hsp90-LA1011 Complex and the Mechanism by which LA1011 may Improve the Prognosis of Alzheimer’s Disease

**DOI:** 10.1101/2023.04.27.538547

**Authors:** S. Mark Roe, Zsolt Török, Andrew McGown, Ibolya Horváth, John Spencer, Tamás Pázmány, László Vigh, Chrisostomos Prodromou

## Abstract

Chaperone systems play a major role in the decline of cognition and contribute to neurological pathologies such as Alzheimer’s disease (AD). While such a decline may occur naturally with age or with stress or trauma, the mechanisms involved have remained elusive. The current models suggest that amyloid-β (Aβ) plaque formation leads to the hyperphosphorylation of tau by a Hsp90 dependent process that triggers tau neurofibrillary tangle formation and neurotoxicity. Several co-chaperones of Hsp90 can influence the phosphorylation of tau, including FKBP51, FKBP52 and PP5. In particular elevated levels of FKBP51 occur with age and stress and are further elevated in AD. Recently, the dihydropyridine, LA1011 was shown to reduce tau pathology and amyloid plaque formation in transgenic AD mice, probably through its interaction with Hsp90 although the precise mode of action is currently unknown. Here, we present a co-crystal structure of LA1011 in complex with a fragment of Hsp90. We show that LA1011 can disrupt binding of FKBP51, which might help to rebalance the Hsp90-FKBP51 chaperone machinery and provide a favourable prognosis towards AD. Clinically, this is highly significant, as AD is generally a disease affecting older patients, where slowing of disease progression could result in AD no longer being life limiting.

## Introduction

Heat shock protein 90 (Hsp90) is a molecular chaperone, which together with its co-chaperones is responsible for the activation of an eclectic set of client proteins [1]. These clients represent key signalling proteins, which when dysregulated lead to disease including cancer and neurodegeneration [2-5]. The role of Hsp90 is well documented in Alzheimer’s disease (AD) [6], which is the most common neurodegenerative disease in humans and is rising due to increases in longevity.

It has been proposed that the overexpression of mutant forms of β-amyloid precursor protein (APP) leads to amyloid-β (Aβ) plaque and tau neurofibrillary tangle formation [5]. Hsp90, forms macromolecular complexes with co-chaperones, which can regulate tau metabolism and Aβ processing [7]. The current model suggests that Aβ peptide triggers the Hsp90 directed hyperphosphorylation of tau (tubulin associated unit), [8,9], that subsequently leads to neurofibrillary tangles and neurotoxicity [10-16]. However, despite significant amounts of research, further clarification of the pathogenetic mechanism is needed.

Abnormal or modified tau can trigger its own degradation by the recruitment of CHIP (Carboxy-terminus of Hsc70-Interacting Protein), an E3 ligase co-chaperone of Hsp90. CHIP drives ubiquitination and subsequently promotes the downstream proteasomal degradation of tau. However, Hsp90 also associates with a variety of other co-chaperones including the peptidylprolyl cis/trans isomerase immunophilins FKBP51 (FK-506-binding protein 51) and FKBP52 and the cyclophilin Cyp40 (Cyclophilin 40 and also known as Peptidylprolyl Isomerase D) [17]. The latter was recently shown to be neuroprotective as it is able to disaggregate tau fibrils *in vitro* and, significantly, it prevented toxic tau accumulation *in vivo* [18]. However, Cyp40 appears to decrease with age and is also repressed in AD [19]. In contrast, FKBP51, is able to preserve toxic tau oligomers *in vivo [20]* and mice lacking FKBP51 were shown to have decreased tau levels in the brain [20,21]. Significantly, FKBP51 levels appear to progressively increase with age and are further increased in AD patients [20,22]. Stress is also known to increase FKBP51 and is associated with AD [23,24]. The structure of FKBP51 in complex with Hsp90 was recently reported and it shows that the seventh helix of the TPR domain of FKBP51 binds across a hydrophobic cleft or pocket formed at the dimer interface at the MEEVD peptide end of the C-terminal domain of Hsp90 [25]. The interaction is critical for aligning the active PPIase domain of FKBP51 with bound tau client.

In contrast, FKBP52, by a direct molecular interaction with truncated and wild-type tau, induces its aggregation [26,27]. However, FKBP52 levels appear to be lower in the cortex of AD patients’ brains [19,28]. Another co-chaperone that plays a role in the phosphorylation state of tau is PP5 (Protein Phosphatase 5). The latter is a Ser/Thr phosphatase that is activated when bound to Hsp90 [29] and is able to dephosphorylate tau at several phosphorylation sites connected to AD pathology [30]. Significantly, PP5 activity is repressed in AD [31]. While the list of co-chaperones that interact with Hsp90 to affect the phosphorylated state of tau may be more extensive, FKBP51, FKBP52 and PP5 appear to be major players and consequently imbalances in these co-chaperones could have major implications in the development of AD [17]. Thus, the maintenance of healthy homeostasis represents a prime focus and is a key therapeutic target for drug discovery with AD and neurodegenerative diseases [32,33]. Consequently, there has been a drive to develop Hsp90 inhibitors that promote the degradation of tau [7,34,35] or alternatively to identify small molecules that can restore the normal balance of chaperone and co-chaperone systems that decline with age or are abnormally altered through stress.

Previously, we found that the dihydropyridine, LA1011, could increase dendritic spine density and reduce tau pathology and amyloid plaque formation in transgenic AD mice [36]. Further work, identified the C-terminal domain of Hsp90 as the target for LA1011 binding and molecular dynamics suggested a specific site to which it bound. Attempts to confirm the site were not wholly conclusive, because mutagenesis of the proposed binding site did not provide a means to prevent binding of LA1011 completely [37]. Nonetheless, these studies identified LA1011 as an inducer of Hsp90 ATPase activity, the heat shock response, and as an effective small molecule against AD in mouse models. Here we now report the crystal structure of a C-terminal domain of Hsp90 in complex with LA1011 and suggest that its binding may help to restore normal levels of FKBP51-Hsp90 complex in cells, which may consequently reduce tau hyperphosphorylation and aggregation. Clinically, this is highly significant as treatment of AD with LA1011 could slow the progression of the disease in elderly patients to a point that AD itself would no longer be life limiting.

## Methods and Materials

### Protein Purification

The yeast Hsp90 C-terminal domain (amino acid residues 438 to 677) was cloned into p3E (A. Oliver, University of Sussex) and expressed in *Escherichia Coli* BL21(DE3) as a GST-tagged fusion. FKBP51 and FKBP51-7He (lacking residues 401-457 of the extended 7^th^ helix of the TPR domain) were a kind gift form D. Southworth (UCSF Weill Institute for Neurosciences) and were expressed with an N-terminal His-tag [25]. The Hsp90 fusion was initially purified using GSH-resin and then cleaved with PreScission protease as previously described [38]. In contrast, FKBP51 constructs were initially purified using Talon affinity metal chromatography [39]. All proteins were further purified using Superdex 200 HR gel-filtration and Q-Sepharose ion-exchange chromatography as previously described [39]. Proteins were dialysed against 20 mM Tris pH 7.4 containing 5 mM NaCl and 1 mM EDTA.

### Crystallization and refinement

The Hsp90-LA1011 complex was crystallized using the sitting-drop method with protein at 22.5 mg ml^−1^ in 20 mM Tris pH 7.4 containing 5 mM NaCl and 1 mM EDTA and 5 mM LA1011 against wells containing 100 mM MES pH 7.5, 200 mM magnesium chloride hexahydrate, 15% PEG Smear Medium (12.5% w/v of each: PEG 3350, 4000, 2000 and 5000 MME) and 5% (V/V) 2-propanol at 14°C (H3 from the BCS Screen HT-96, Molecular Dimensions, MD1-105). Crystals were harvested by successive transfer into a crystallization buffer with increasing ethylene glycol content to 30% and then flash-cooled in liquid nitrogen. Diffraction data were collected from crystals cooled to 100 K on I24 at Diamond Light Source. Data were automatically processed with autoPROC/STARANISO (STARANISO available at http://staraniso.globalphasing.org/cgi-bin/staraniso.cgi, Cambridge, United Kingdom: Global Phasing Ltd.) [40] and the asymmetric unit contents were estimated using the Matthews coefficient (*CCP*4 suite) [41]. The LA1011-Hsp90 structure was solved using *Phaser* (*CCP*4) with a model created from PDB 2CGE. The structure was refined with *REFMAC* 5.7 [42] and manual rebuilding was performed in *Coot* [43]. The structure was deposited in the PDB database and was displayed using *PyMOL* (Schrödinger, L. & DeLano, W., 2020. *PyMOL*, Available at: http://www.pymol.org/pymol).

### Isothermal Titration calorimetry

Heat of interaction was measured on an ITC_200_ microcalorimeter (Microcal) under the same buffer conditions (20 mM Tris, pH 7.5, containing 5 mM NaCl). Aliquots of 1 mM LA1011 were injected into 30 μM of yeast Hsp90 or Hsp90-FKBP51 complex (consisting of 30 μM Hsp90 and 60 μM FKBP51 or FKBP51-7He) at 30°C. Interactions with FKBP51 and FKBP51-7He were performed by injecting aliquots of 400 μM of intact FKBP51 or FKBP51-7He construct into 30 μM of Hsp90 or 30 μM Hsp90 in complex with 1 mM LA1011. Heats of dilution were determined by diluting injectant into buffer. Data were fitted using a curve-fitting algorithm (Microcal Origin).

## Results and Conclusions

### The crystal structure of Hsp90-LA1011

The structure of a complex between LA1011 (Fig. 1A) and a Hsp90 fragment (residues 438-677) was solved by molecular replacement and refined to an anisotropic resolution limit of 2.94 A (isotropic 3.18 Å) (Table 1). The structure reveals two dimer molecules of the Hsp90 fragment in the asymmetric unit, but only one of the dimers contained bound LA1011. LA1011, was found to bind in a small hydrophobic pocket at the extreme C-terminus of the dimer interface of Hsp90 (Fig. 1B-D). This is in contrast to, and at the opposite end of the C-terminal domain, to the previous binding prediction by molecular dynamics [37]. However, one of the methyl ester groups of LA1011 binds across the two-fold symmetry point of the Hsp90 dimer and hence only a single molecule of LA1011 is able to bind. This is in agreement with previous binding studies using isothermal titration calorimetry [37]. The binding pocket is lined by a series of hydrophobic and hydrophilic amino acid residues, but there are no obvious hydrophilic interactions (Fig. 1C and D). The pocket is lined by the amino acids Leu 671, Ile 672, Leu 674, Gly 675, Leu 676 form one molecule of the dimer and by Glu 624, Leu 625, Arg 628, Thr 638, Asp 641, Leu 642, Leu 671 and Leu 674 form the other protomer.

**Table 1.** Crystallographic Statistics

| **Data Set**<br>**(Highest shell in**<br>**parentheses)** | **AutoPROC** | **STARANISO** |
| --- | --- | --- |
| a (Å) | 97.71 |  |
| b (Å) | 97.71 |  |
| c (Å) | 275.57 |  |
| $\alpha$ (°) | 90.0 | |
| $\beta$ (°) | 90.0 | |
| $\gamma$ (°) | 90.0 | |
| Space Group | P 4<sub>1</sub> 2<sub>1</sub> 2<sub>1</sub> |  |
| Wavelength (Å) | 0.96864 |  |
| Resolution Limit (Å) | 92.1–3.18 (3.23–3.18) | 92.1–2.94 (3.19–2.94) |
| Number of Unique Obs. | 22841 (1103) | 23054 (1153) |
| Completeness (%) | 97.4 (97.0) | 91.9 (58.1) |
| Multiplicity | 9.3 (5.9) | 9.2 (4.0) |
| Rmerge (I) % | 0.218 (1.867) | 0.217 (1.215) |
| Rpim(I) % | 0.073 (0.803) | 0.073 (0.666) |
| CC<sub>1/2</sub> | 0.995 (0.408) | 0.995 (0.361) |
| I/σI | 9.0 (1.1) | 9.0 (1.4) |
| Refinement |  |  |
| Resolution Range (Å) |  | 79.7-2.94 |
| Rcryst |  | 0.2470 |
| Rfree |  | 0.3062 |
| Number of protein atoms |  | 6816 |
| Number of ligand atoms |  | 0 |
| Number of solvent atoms |  | 1 |
| Mean B |  | 70.11 |
| Rmsd bond lengths (Å) |  | 0.005 |
| Rmsd bond angles (°) |  | 1.051 |

**Figure 1.**
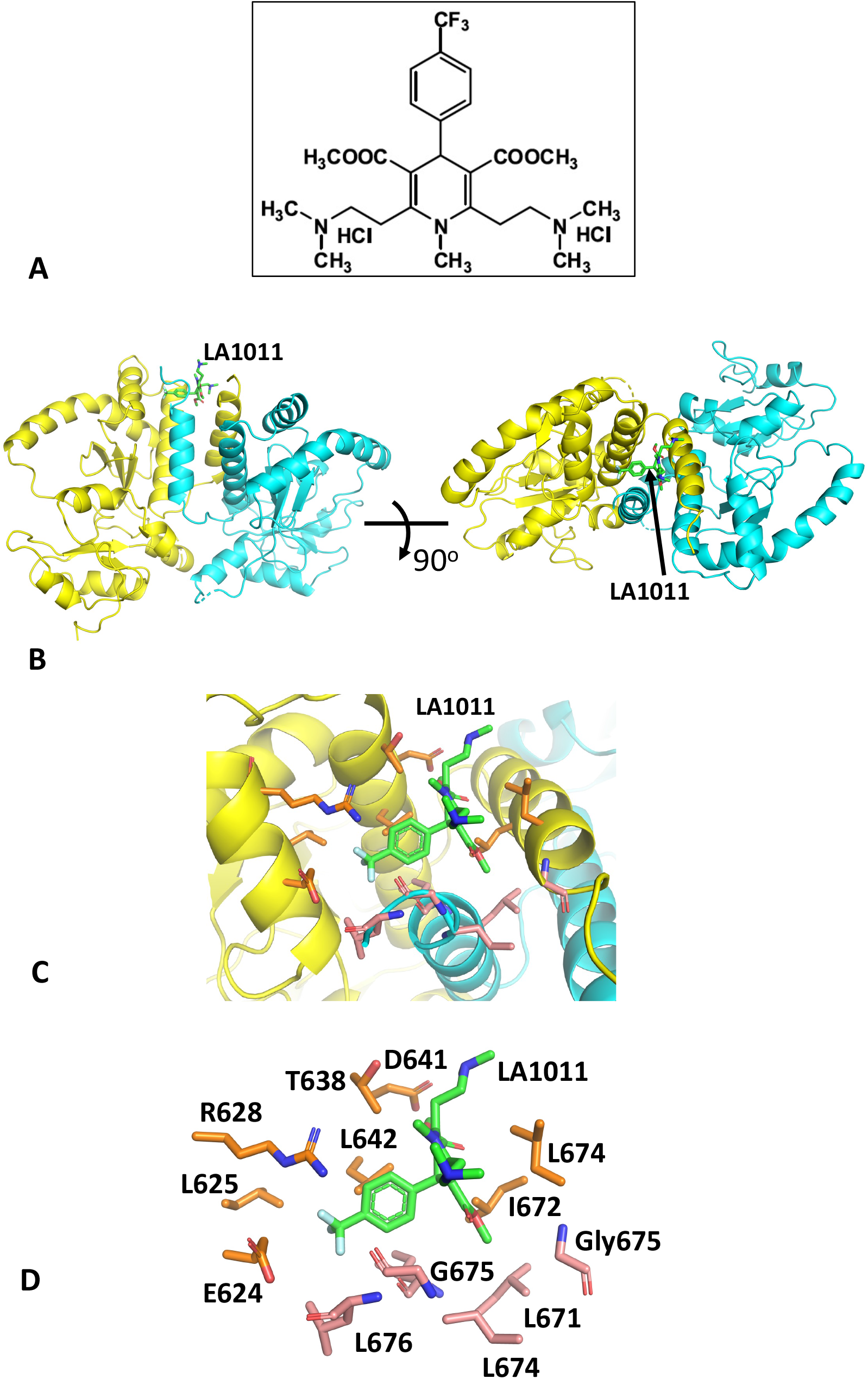
The structure of the Hsp90-LA1011 complex. A), Structure of LA1011. B), PyMol cartoon showing orthogonal views of the Hsp90-LA1011 complex. The two protomers of the C-terminal domain of the Hsp90 dimer are shown in yellow and cyan. LA1011 is shown in green stick format. C), PyMol cartoon showing a close up of the LA1011 binding site of Hsp90. The two protomers of the C-terminal domain of the Hsp90 dimer are shown in yellow and cyan and amino acid side-chains in gold and salmon stick format, respectively. LA1011 is shown in green stick format. D). PyMol cartoon showing the side chains of Hsp90 (gold sticks form protomer A and salmon sticks form protomer B) that form the hydrophobic pocket of Hsp90 to which LA1011 (green sticks) is bound.

### LA1011 and FKBP51 competitively bind to Hsp90

The structure of Hsp90 in complex with FKBP51 has previously been determined [25], which shows that helix 7 of the TPR domain of FKBP51 is docked in a pocket or cleft formed at the extreme C-terminus of the Hsp90 dimer interface (Fig. 2). Previously, molecular dynamics had predicted that LA1011 bound to a similar pocket at the other end of the C-terminal dimer interface [37]. However, mutagenesis could not prove beyond any doubt that this site was the actual binding site observed in interaction studies, because there was a lack of residues that could be mutated to sterically hinder binding altogether. However, we wondered whether the true binding site could be the same as that seen for helix 7 of the TPR domain of FKBP51. We therefore decided to use isothermal titration calorimetry (ITC) to investigate whether FKBP51 binding could block LA1011’s interaction with Hsp90 and *vice versa*.

**Figure 2.**
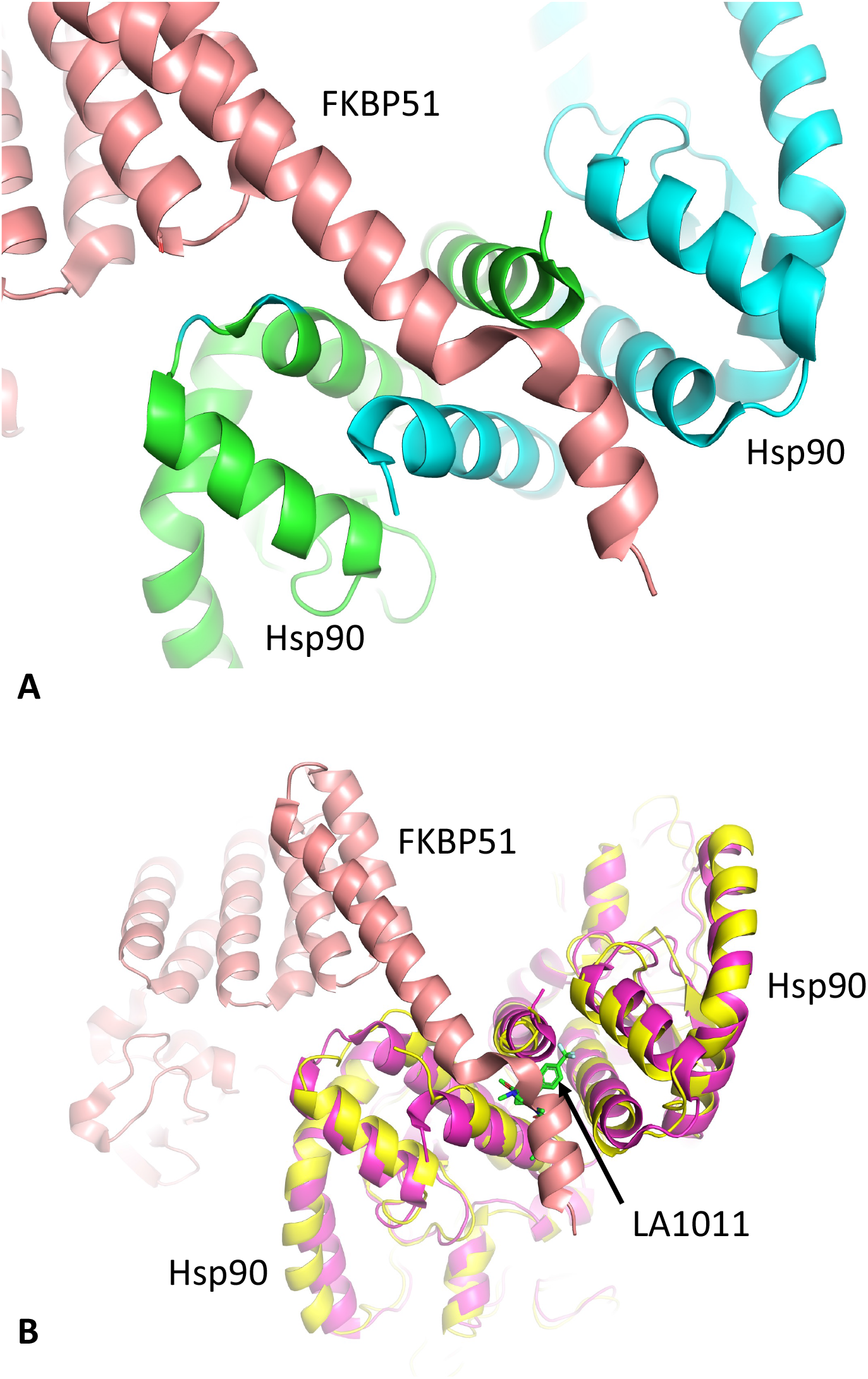
The structure of the Hsp90-FKBP complex. A), PyMol cartoon showing helix 7 (salmon) of the FKBP51 TPR-domain binding to the C-terminal domains of Hsp90 (green and cyan). B), Superimposition of the Hsp90-FKBP51 complex (magenta and salmon, respectively) with the Hsp90-LA1011 complex (yellow and green sticks, respectively). Competition for binding between LA1011 and helix 7 of FKBP51 to the extreme C-terminal hydrophobic pocket of Hsp90 is evident through steric clashes with LA1011.

Since only one molecule of FKBP51 is able to interact with the extreme C-terminal hydrophobic pocket of Hsp90, but two Hsp90 MEEVD peptide binding sites are available, we expected that two different binding events should be detectable by ITC for intact FKBP51, but a single-site would be a better fit for the FKBP51 mutant lacking the helical extension (amino acid residues 401-457; FKBP51-7He) of the TPR domain. As expected, full-length FKBP51 bound Hsp90 and we found that the one-site model fitting of the data was poor compared to a two-site model fit (Fig. 3A and B). The *K*d for the two-site binding was 0.5 and 32.4 μM (Fig. 3B), indicating that the C-terminal extension of the TPR domain of FKBP51 was an essential component for high affinity binding to Hsp90. We next measured the affinity of binding for FKBP51-7He with Hsp90, which we expected to produce data that would conform to a one-site model fit, since two molecules of the construct would have equivalent binding sites (MEEVD only). As expected, using a one-site model fit, we found that this construct bound with a *K*d of 11.6 μM and with a stoichiometry (1:1.25, Hsp90:FKBP51-7He), that was close to one (Fig. 3C). The affinity for this interaction was similar to the weaker binding affinity obtained using the two-site model fit with intact FBBP51, which is due to MEEVD binding alone (32.4 μM; Fig. 3B). This indicates, as we had expected, that high affinity binding to Hsp90 had been compromised with the FKBP51-7He mutant. We next measured the binding affinity of our most recent sample of LA1011 and found it to bind Hsp90 with a *K*d of 13.1 μM (Fig. 3D), which was similar to our previous studies (*K*d = 13.5 μM) [37]. To investigate the binding of LA1011 to a Hsp90-FKBP51 complex, we used a 30 μM Hsp90 solution saturated with 60 μM of intact FKBP51. We found that the binding of LA1011 to the Hsp90-FKBP51 complex was compromised (*K*d = 108 μM; Fig. 3E), but restored when using Hsp90 saturated with the FKBP51-7He mutant (*K*d = 17.9 μM; Fig. 3F). This affinity, for the binding of LA1011 to Hsp90-FKBP51-7He, was similar to that for the binding of LA1011 to free Hsp90 (*K*d = 13.1 μM; Fig. 3D). The results so far suggest that the helical extension of the TPR domain of FKBP51 and LA1011 compete for binding to the extreme C-terminal hydrophobic pocket of Hsp90.

**Figure 3.**
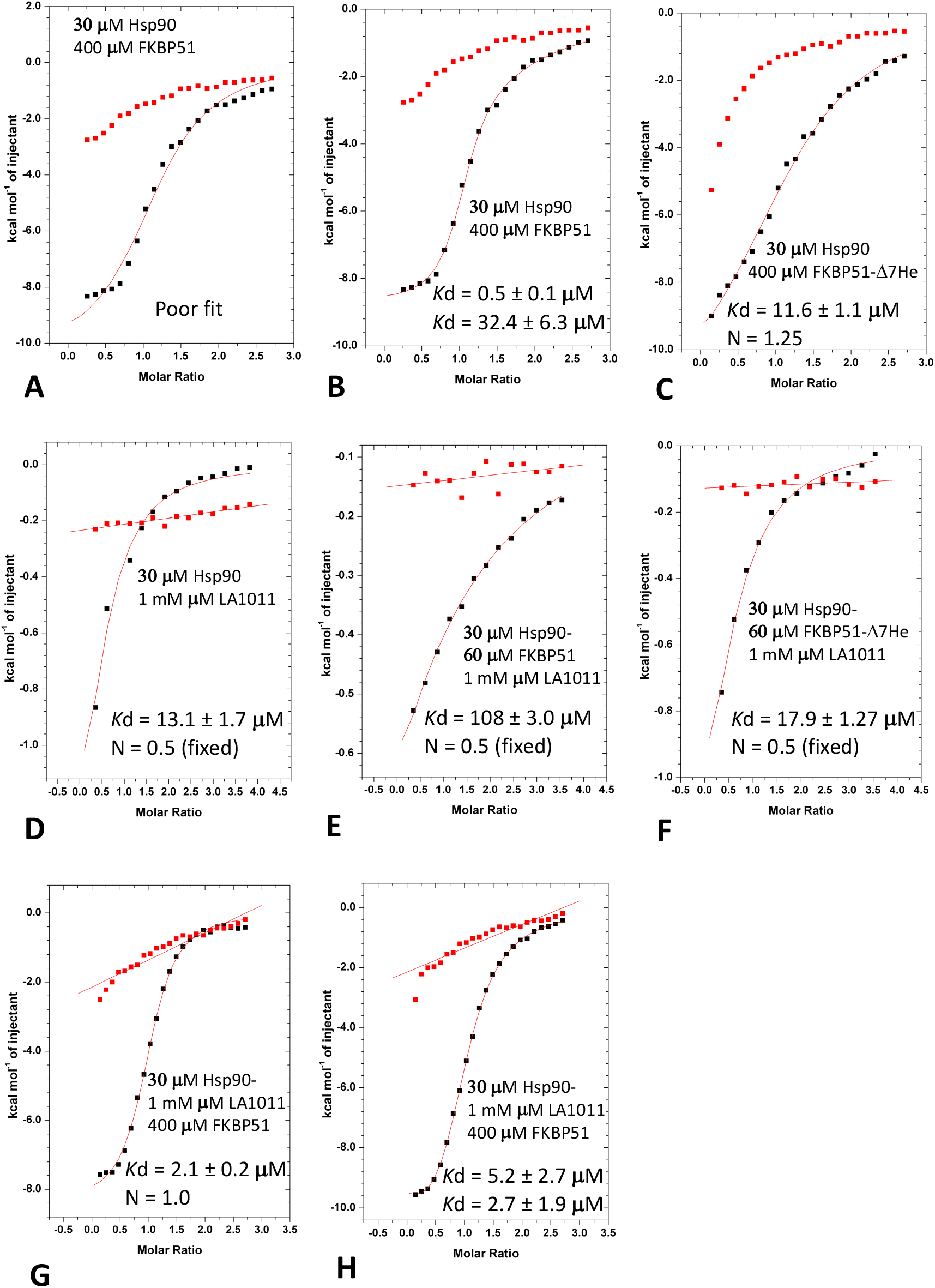
Isothermal titration calorimetry binding studies for FKBP51 and LA1011 with Hsp90. A) Titration of FKBP51 into Hsp90 and fitted with a one-site binding model that does not fit well to the experimental data. B), Titration of FKBP51 into Hsp90 and fitted as a two-site binding model, which fits the experimental data better than a one-site binding model (Fig 3A). C), Binding of the FKBP51-ΔHe construct to Hsp90 showing that the experimental data can be fitted with a one-site binding model. D), Binding of LA1011 to Hsp90. E), Binding of LA1011 to the Hsp90-FKBP51 complex and F), to the Hsp90-FKBP51-7He complex, showing that LA1011 binding is only compromised with the Hsp90-FKBP51 full-length complex. G), Binding of FKBP51 to the Hsp90-LA1011 complex fitted as a one-site binding model and H), fitted as a two-site binding model. The affinities in both fittings for binding of FKBP51 are all similar and do not show the high affinity binding observed in Fig 3 B (Kd = 0.5 μM), suggesting that FKBP51 only binds to the MEEVD motif of Hsp90 in the presence of LA1011.

To further validate this conclusion, we next investigated how LA1011 affected the binding of intact FKBP51. We therefore formed a complex consisting of 30 μM Hsp90 and 1 mM LA1011 and measured the binding for intact FKBP51. Using a one-site binding fit for intact FKBP51 gave an affinity for binding of *K*d = 2.1 μM (Fig. 3G), while a two-site fit gave affinities of *K*d = 5.2 and *K*d = 2.7 μM (Fig. 3H). Neither fit could reproduce the high affinity binding of intact FKBP51 to Hsp90 (Kd = 0.5 μM; Fig. 3B) previously seen, suggesting that binding of intact FKBP51 to the extreme hydrophobic pocket of Hsp90 had been compromised. The results, collectively, show that only one molecule of intact FKBP51 simultaneously binds the MEEVD motif and the hydrophobic pocket at the extreme C-terminal dimer interface of Hsp90 and that binding of a second molecule is limited to just the MEEVD motif in our *in vitro* system. However, when the hydrophobic pocket of Hsp90 is occupied by LA1011, FKBP51 molecules are limited to binding just the MEEVD motif of Hsp90.

### Effects of LA1011 on the binding of other TPR domain containing co-chaperones of Hsp90

The binding of LA1011 to the extreme C-terminal hydrophobic pocket of Hsp90 clearly disrupts the binding of FKBP51. However, other TPR domain containing co-chaperones also contain an equivalent extended helix 7 of their TPR domains, which may also bind the C-terminal hydrophobic pocket of Hsp90. The most important residues for the binding of the helical extension of FKBP51 are Ile 408, Tyr 409, Met 412, Phe 413, Phe 416 and Ala 417 (IY--MF--FA) (Fig. 4 A,B). FKBP52 has a similar conserved motif, (LY--MF--LA), where Ile 408 and Phe 416 of FKBP51 are replaced by Leu 409 and Leu 417 in FKBP52, respectively (Fig. 4B). The degree of conservation suggests that both co-chaperones would bind in a similar manner to the extreme C-terminal hydrophobic pocket of Hsp90. Cyp40 on the other hand appears to have what looks like a truncated motif (VY--MF-), where Ile 408 is replaced by Val 364 and the downstream residues, equivalent to Phe 416 and Ala 417 of FKBP51, are missing altogether in Cyp40 (Fig. 4B). Another example of a TPR helix extension binding to the hydrophobic pocket of Hsp90 was recently seen in two EM structures of Hsp90 in complex with PP5 (Serine/threonine-protein phosphatase), CDC37 and Kinase [44,45]. Significantly, it was reported that Phe 148 and Ile 152 of human PP5 is bound in the extreme C-terminal hydrophobic pocket of Hsp90 [45], but our structural comparisons show that helix 7 of the TPR domain adopts a completely different conformation relative to that of FKBP51 (Fig. 4C). The TPR helical extension of human PP5 also possess two conserved acidic residues, Asp 155 and Glu 156, that are in a close proximity to basic residues, Arg 679 and Arg 682, of Hsp90 [45]. In yeast PPT1, these residues are also conserved (Phe 132 (PP5-Phe 148), Ile 136 (PP5-Ile 152) and Glu 140 (PP5-Glu156) except for Asp 155 of human PP5 which is an alanine residue instead (Ala 139). The interaction of the TPR helical extension of PP5 with the hydrophobic pocket of Hsp90, consequently, does not seem to have a similarly conserved FKBP51-like binding motif (Fig. 4B). Nonetheless, the binding of the helical extension of PP5 would most likely also be compromised by Hsp90 bound LA1011 (Fig. 4D). In another study, mutation of the TPR helical extension of AIP (aryl hydrocarbon receptor-interacting protein) was found to be important for client protein maturation [38]. Consequently, the mechanism by which AIP-dependant client maturation could be affected might be because of a disruption of AIP helical extension docking with the extreme C-terminal hydrophobic pocket of Hsp90. Thus, failure to correctly dock would presumably compromise client protein maturation. We note that human AIP has two conserved phenylalanine residues (Phe 324 and 328), which align with the conserved Phe 148 and Ile 152 residues of PP5 (Fig. 4B).

**Figure 4.**
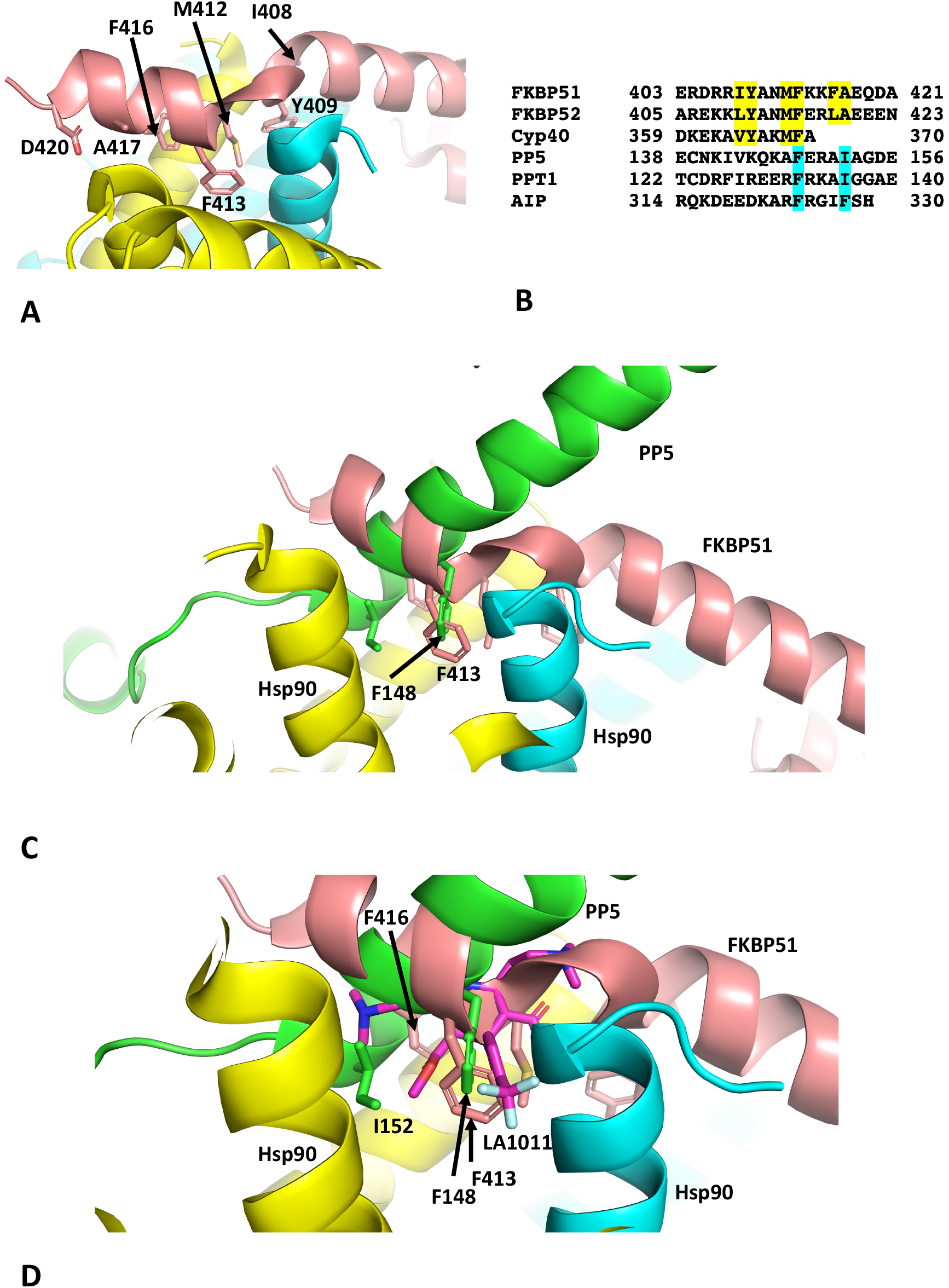
TPR domain binding to the extreme C-terminal hydrophobic pocket of Hsp90. A), PyMol cartoon showing the critical residues (stick format) of FKBP51 (salmon) that interact with the hydrophobic pocket of Hsp90 (yellow and cyan). B), Sequence alignment of segments of the helix extension of selected TPR domain containing co-chaperones of Hsp90. Amino acid numbering is shown. Yellow highlight, conserved residue positions for FKBP51, FKBP52 and Cyp40; cyan, the conserved residues (Phe 148 and Ile 152 in human PP5; Phe 132 and Ile 136 in *S. cerevisiae* PPT1 and Phe 324 and 328 in human AIP) involved in binding to the hydrophobic pocket of Hsp90. C), PyMol cartoon comparing the conformation of the bound TPR helix extension of FKBP51 (salmon) and that of PP5 (green) to the hydrophobic pocket of Hsp90 (cyan and yellow). D) PyMol cartoon showing that LA1011 (magenta sticks) sterically clashes with the TPR helix extension of PP5 (green) and FKBP51 (salmon). Yellow and cyan are Hsp90.

This brief analysis suggests that the docking conformation of the helical extension of TPR domain containing co-chaperones of Hsp90 might be variable or at least there are two conformational binding states so far observed. In both these states a key phenylalanine residue is well placed in the binding position of the trifluoromethyl phenyl group of LA1011 (Fig. 4D). However, the effects of LA1011 on the functioning of TPR domain containing co-chaperones of Hsp90 may be selective and those TPR domains that might not dock with the C-terminal hydrophobic pocket of Hsp90 could naturally be unaffected. Clearly, further work on this is required to establish a clearer picture.

## Discussion

Previous results using molecular dynamics suggested that LA1011 bound to the C-terminal domain of Hsp90, but mutagenesis could not univocally confirm the exact site of interaction. We therefore set out to obtain a crystal structure of the Hsp90-LA1011 complex to determine the precise binding site. The crystals of the complex showed that LA1011 binds to the opposite end of the C-terminal domain of Hsp90 in relation to the site suggested by molecular dynamics. However, some caution should be exercised in completely disregarding the molecular dynamics site, as both sites could be targets of LA1011, but that our crystal structure is selective for the extreme C-terminal hydrophobic pocket. We in fact note, that the binding of LA1011 to a Hsp90-FKBP51 complex does not in itself completely block LA101 binding. This could be due to competition between FKBP51 and LA1011, or alternatively binding of LA1011 could be occurring at more than one site, such as the molecular dynamics site identified previously. It is clear that further work is required in order to clarify the exact situation. However, this does not escape the fact that it is clear that LA1011 directly competes with FKBP51 for binding to the extreme C-terminal hydrophobic pocket of Hsp90. It appears that the trifluoromethyl phenyl group of LA1011 to some degree mimics a key binding residue of FKBP51 (Phe 413), but also that of PP5 (Phe 148) and perhaps other co-chaperones, such as AIP (Phe 324). The binding interactions and conformations of FKBP51 and PP5 to the C-terminal hydrophobic pocket of Hsp90, will now form the basis for the design of improved small molecules that can interact with this Hsp90 hydrophobic pocket, which we are now actively pursuing.

Previous studies suggest that FKBP51 is a major player in AD, as it increases with age, with stress, increases even further with AD disease and significantly it increases the hyperphosphorylation of tau [20,22]. Since FKBP51 is thought to favour tau fibrillation, it would be reasonable to assume that decreasing the association of FKBP51 with tau bound Hsp90 would decrease the ability of this system to promote the hyperphosphorylation of tau [8,9]. We therefore suggest that our previous observations where LA1011 increased dendritic spine density and reduced tau pathology and amyloid plaque formation in transgenic AD mice, could be directly due to the disruption of the Hsp90-FKBP51 complex [36]. Thus, *in vivo* where the cellular concentration of the Hsp90-FKBP51 complex is ‘abnormally’ elevated (due to high levels of FKBP51), LA1011 might restore ‘normal’ cellular levels of the Hsp90-FKBP51 complex by reducing the binding of FKBP51 to Hsp90. This in turn could then reduce the phosphorylation of tau and might therefore limit disease progression. Further biological studies using cells and a mouse model of AD are in progress to gain direct evidence on our concepts described herein.

Another co-chaperone found to associate with the extreme C-terminal hydrophobic pocket of Hsp90 is the serine-threonine tyrosine kinase PP5. However, the mode of binding is different to that reported for FKBP51 [25,44]. It also appears likely that AIP may also interact in a similar way due to client protein dependency on the extended helix of its TPR domain, but further work is required to conclusively establish this [38]. Disruption of the interaction of AIP with Hsp90 may be a mechanism sufficient for pituitary adenoma predisposition [38]. However, the fact that numerous TPR domain containing co-chaperones of Hsp90 are likely to be affected by LA1011, it does suggest that numerous Hsp90 processes could be affected by targeting the extreme C-terminal hydrophobic pocket of Hsp90. Nonetheless, the mouse model studies suggest a positive prognosis with LA1011 treatment [36] and consequently this makes this target of prime importance for further studies, especially as drugs to treat AD have been difficult to develop due to a lack of understanding of the underlying mechanism that leads to this disease [46]. Here we present a possible mechanism that appears to influence AD progression that could be targeted with a small molecule to help improve the prognosis of AD. Since AD is generally a disease that affects the elderly, small changes in the progression of such a disease could have profound effects on the quality of life of such patients.

## Author contributions

Conceptualization, C.P.; methodology, S.M.R., A.G., Z.T and C.P.; formal analysis, C.P and S.M.R.; writing—original draft, C.P.; writing—review & editing, C.P., I.H., J.S, T.P., A.M., and L.V.; supervision, C.P, I.H., J.S, T.P., and L.V. All authors have read and agreed to the published version of the manuscript.

## Institutional Review Board Statement

Not applicable.

## Informed Consent Statement

Not applicable.

## Data Availability Statement

Raw data for ITC experiments can be found at https://doi.org/10.25377/sussex.22341712. The structure was deposited in the PDB database.

## Acknowledgments

The owner of the patent application associated with the research compound, LA1011, described herein is Richter Gedeon Plc. pharmaceutical company Patent number 10660789 (https://patents.justia.com/patent/10660789).

